# Native protein delivery into rice callus using ionic complexes of protein and cell-penetrating peptides

**DOI:** 10.1101/570853

**Authors:** Boyang Guo, Jun Itami, Kazusato Oikawa, Yoko Motoda, Takanori Kigawa, Keiji Numata

**Affiliations:** Biomacromolecules Research Team, RIKEN Center for Sustainable Resource Science, Japan; Laboratory for Cellular Structural Biology, RIKEN Center for Biosystems Dynamics Research, Japan

**Author notes:** Corresponding author Keiji Numata, Ph.D. Biomacromolecules Research Team, RIKEN Center for Sustainable Resource Science, 2-1 Hirosawa, Wako-shi, Saitama 351-0198, Japan.

## Abstract

Direct protein delivery into intact plants remains a challenge for the agricultural and plant science fields. Cell-penetrating peptide (CPP)-mediated protein delivery requires the binding of CPPs to a protein to carry the protein into the cell through the cell wall and lipid bilayer. Thus, we prepared ionic complexes of a CPP-containing carrier peptide and a cargo protein, namely, Citrine yellow fluorescent protein, and subsequently studied their physicochemical properties. Two types of carrier peptides, BP100(KH)_9_ and BP100CH_7_, were investigated for delivery efficiency into rice callus. Both BP100(KH)_9_ and BP100CH_7_ successfully introduced Citrine protein into rice callus cells under pressure and vacuum treatment. Moreover, delivery efficiency varied at different growth stages of rice callus; 5-day rice callus was a more efficient recipient for Citrine than 21-day callus.

## Introduction

With the rapid growth of the global population, biotechnological breeding to improve the quality and quantity of rice, a major agricultural crop, has attracted a great deal of attention. For example, CRISPR/Cas9 transgenic technology has been widely used in rice cells to introduce genome modifications [1-5] based on the Agrobacterium-mediated [1, 2, 5] and protoplast transformation methods [3, 4]. However, these processes have a strong possibility of causing off-target mutations [6], which leads to a safety issue in major crops, namely, damage to ecological equilibrium [7]. Previously, it was reported that direct protein delivery using a carrier could avoid the risk of incorporating exogenous genes into the genome through nuclear transformation [8, 9]. Thus, the direct introduction of proteins into plant cells is of great interest for various application fields, including plant genome editing.

Cell-penetrating peptides (CPPs), also called protein transduction domains, are short peptides that facilitate the transport of cargo molecules through the plasma membrane to the cytosol [10]. In many cases, CPPs are coupled to cargo molecules through covalent conjugation, forming CPP-cargo complexes [11, 12]. To date, DNA, RNA, nanomaterials and proteins such as antibodies have been reported as cargo molecules that can be combined with CPP [9–12]. Most studies of CPP-protein complexes have contributed to applications in mammalian cells, whereas few studies have focused on plant cells. This is because native proteins are large molecules with specific folding structures, unlike nucleic acids, nanomaterials and antibodies. Additionally, the surface charges of proteins differ from one another. Notably, plant cells are surrounded by the cell wall, which prevents the internalization of large molecules [9, 13]. The plant cell wall mainly contains cellulose, hemicellulose and pectin [13], and its biochemical composition changes during plant growth [14]. Thus, we need to optimize various conditions to achieve delivery of native proteins into plant cells through the cell wall.

BP100(KH)_9_ (amino acid sequence: KKLFKKILKYLKHKHKHKHKHKHKHKHKH) is a fusion peptide containing CPP and cationic sequences (**Fig. 1**). It was previously demonstrated that BP100(KH)_9_ is able to transport native proteins in the size range of 27 to 160 kDa into *Arabidopsis thaliana* leaf cells [15]. This protein delivery system explored the possibility of penetrating plant cells with native proteins for biotechnological plant modification. Another CPP-containing peptide for plant cells is BP100CH_7_ (amino acid sequence: KKLFKKILKYLHHCRGHTVHSHHHCIR) (**Fig. 1**), which was designed as a stimulus-response peptide and could release DNA into the cytoplasm after cellular uptake [16]. In this study, to achieve delivery of complexes of CPPs and native proteins into rice callus, we adopted these two types of CPP-containing fusion peptides (CPP-FPs), BP100(KH)_9_ and BP100CH_7_, to prepare ionic complexes with a yellow fluorescent protein, Citrine (**Fig. 1**). After Citrine is introduced into the cell by the CPP-FP, its fluorescence signals can be detected by confocal laser scanning microscopy (CLSM). Furthermore, to determine an efficient protein delivery condition using the CPP-FP complex, rice callus cells in different growth stages were evaluated as target plant materials. Based on these results, we determined the best combination of CPP-FP/Citrine complex and recipient rice callus cell.

**Figure 1.**
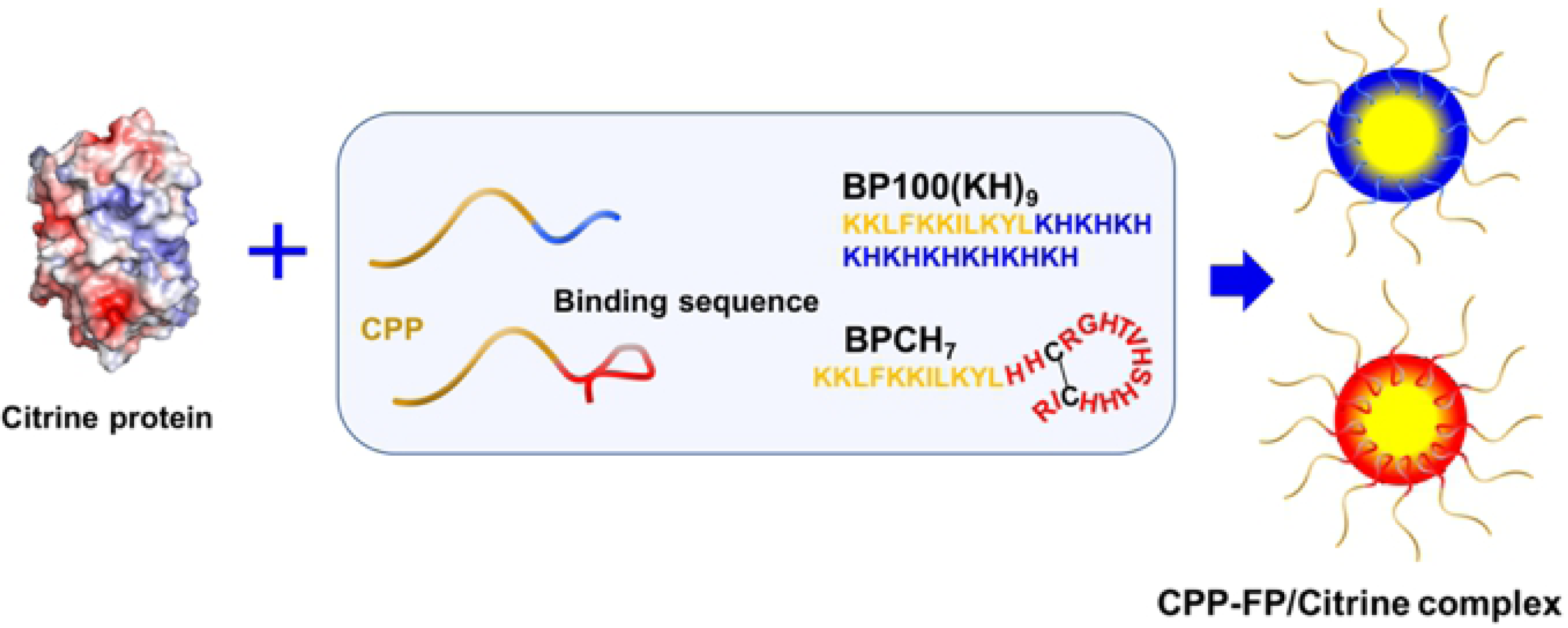
Schematic diagram of the peptide-based protein delivery system. The negatively charged Citrine was ionically combined with the molecular binding sequence of the designed peptides to form the CPP-FP/Citrine complexes.

## Materials and Methods

### Plant Growth Conditions, Embryogenic Callus Induction and Recipient Cells

Mature dry seeds of *Oryza sativa* cv. Nipponbare (*O. sativa*) were surface-sterilized with 70% ethanol (v/v) for 1 min, followed by 30 min in 50% (v/v) commercial bleach with rotation at 20 rpm. The seeds were then washed 8-10 times with sterile distilled water and dried on autoclaved 3 mm Kimwipes (Kimwipes S-200, Nippon Paper Crecia Co., Ltd.) for 5 min. For callus induction, sixteen seeds were inoculated on callus induction medium (N6D) and incubated at 30°C with continuous light in a plant bioincubator (CLE-303 cultivation chamber, TOMY Seiko Co. Ltd., Tokyo, Japan). N6D was prepared using basal 30 g/L lactose, 0.3 g/L casamino acid, 2.8 g/L L-proline, 2 mg/L 2,4- dichlorophenoxyacetic acid (2,4-D), 4.0 g/L Chu (N_6_) basal salt mix, and 4 g/L Phytagel. The pH was adjusted to 5.8 before autoclaving. After 5 days of cultivation, the callus was cut into approximately four equal parts. This callus was used as the 5-day callus in this study. Then, after 21 days of cultivation, the self-shedding callus was collected and used as the 21-day callus. For the callus regeneration test, the callus was cut into small pieces of different sizes (approximately 0.5 to 6.0 mm), transferred onto a regeneration medium and incubated at 30°C with continuous light for 7 days. The regeneration medium was prepared using 4 g/L MS powder with vitamins, 30 g/L sorbitol, 30 g/L sucrose, 4 g/L casamino acid, 2 mg/L 2,4-D, 2 mg/L 1-naphthaleneacetic acid (NAA), and 4 g/L Phytagel, and the pH was adjusted to 5.8 before autoclaving. Callus that generated green plants was considered to possess regeneration ability. The chemicals used in this research were purchased from Sigma (Sigma-Aldrich, MO, USA) and Wako (Fujifilm Wako Pure Chemical Corporation, Osaka, Japan).

### Preparation of the CPP-FP/Citrine Complexes

The CPP-FPs used in this study were synthesized by the solid-phase method and purified as described previously [17]. The Citrine protein was prepared by cell-free synthesis and purified as reported previously [18, 19]. To prepare the CPP-FP/Citrine complexes, 2 μg Citrine (1 mg/mL) was mixed with CPP-FP (1 mg/mL) at various molar ratios. The BP100(KH)_9_/Citrine complex was prepared at molar ratios of 1, 5, 10, 20 and 30, whereas the BP100CH_7_/Citrine complex was prepared at molar ratios of 1, 5, 10, 20, 30, 50, and 100. The complex solutions were pipetted gently and incubated at 25°C for 30 min under dark conditions. This solution was adjusted to the final volume of 100 μL by adding autoclaved Milli-Q water and then continuing the incubation under the same conditions for another 30 min. After 10- fold dilution, each solution was characterized by Zetasizer Nano ZS (Malvern Instruments Co. Ltd., Worcestershire, UK) to analyze the size, polydispersity index (PDI) and zeta potential of the complexes, as previously reported [15].

### Penetration of CPP-FP/Citrine Complexes into Rice Callus Cells

Samples of 5-day or 21-day rice callus (10 mg) were immersed in 100 μL of each complex solutions in a screw cap tube. Subsequently, the rice callus in the complex solution was depressurized at −0.08 MPa for 1 min and compressed at +0.08 MPa for 1 min. After this treatment, the callus was transferred onto N6D medium and incubated at 30°C under dark conditions until use.

### Intracellular Uptake and Distribution Analysis of CPP-FP/Citrine Complexes

CLSM (Zeiss LSM 700, Carl Zeiss, Oberkochen, Germany) was used to evaluate the cellular uptake of the CPP-FP/Citrine complex every 24 hours. Before observation by CLSM, the callus on N6D was transferred into a 1.5 mL Eppendorf tube and washed thoroughly with Milli-Q water containing 0.1% TWEEN 20 (Sigma-Aldrich) five times to remove excess Citrine on the cell surface. Thereafter, the callus was cut into tiny pieces (0.5 mm in diameter), mounted on glass slides, and covered with a coverslip. The CPP-FP/Citrine complex was detected by setting the excitation to 488 nm and emission to a range of 505-600 nm. To distinguish the plasma membrane region and cytoplasm region, the callus was additionally incubated with FM4-64 (20 μM, 20 min at 25°C) for cell membrane staining; this fluorophore was detected by setting the excitation to 405 nm and emission to 560–700 nm. Furthermore, quantification of intracellular Citrine was performed by Western blot immunoassay. To extract the protein from the rice callus cells at 72 hours postinfiltration, the rice callus was frozen with liquid nitrogen and then ground into powder in a mortar. Fifty microliters of 10 mM Tris-HCl buffer (pH 7.4) containing 10 μL Halt protease inhibitor cocktail (Thermo Fisher Scientific, Waltham, MA) was added to 100 mg of callus powder, mixed and incubated on ice for 1 hour. After centrifugation of the sample at 150 rpm and 4°C for 20 min, the supernatant was collected and used as the cell extract. Ten microliters of the cell extract was subjected to 4-20% sodium dodecyl sulfate polyacrylamide electrophoresis gels (SDS-PAGE, Bio-Rad, CA). SDS-PAGE containing proteins were transferred onto PVDF membranes (Bio-Rad) using a semidry transfer cell (Bio-Rad) at 10 V for 1 hour. The immunodetection of Citrine was performed with a rabbit polyclonal antibody (GFP antibody NB600-308, Novus Biologicals Co., dilution 1:2000). IgG goat anti-rabbit antibody conjugated with horseradish peroxidase was used as a secondary antibody (goat anti-rabbit IgG H&L HRP ab6789, Abcam, Cambridge, UK, dilution 1:20000). A Luminescent Image Analyzer (LAS-3000, Fujifilm, Tokyo, Japan) was used to visualize the Citrine.

## Results

### Physicochemical Characterization of the CPP-FP/Citrine Complexes

To examine the effects of the CPP-FP/Citrine molar ratio (P/C ratio) in forming the ionic complex, we first prepared complexes of CPP-FP/Citrine in a range of 1 to 30. The results revealed that, without CPP-FP decoration, the hydrodynamic diameter of Citrine was approximately 240 nm (**Fig. 2a-b**). The hydrodynamic diameter and PDI of the BP100(KH)_9_/Citrine complex decreased at P/C ratios from 1 to 10 and increased at P/C ratios from 10 to 30. At a P/C ratio of 1, the complex showed a larger average diameter and PDI value (**Fig. 2a, Table S1**), suggesting that these CPP-FP/Citrine complexes were more aggregates than ordered micelles. By increasing the P/C ratio, the complex hydrodynamic diameters and PDI of BP100(KH)_9_/Citrine became uniform and homogeneously distributed at a P/C ratio of 10 (**Fig. 2a, Table S1**). After the addition of BP100(KH)_9_, the zeta potentials of the complexes changed from negative to positive at all molar ratios of the BP100(KH)_9_/Citrine complex (**Fig. 2a**). In contrast, BP100CH_7_/Citrine at various P/C ratios did not show a significant difference in diameter and PDI, but a more negative zeta potential was observed at higher P/C ratios (**Fig. 2b, Table S1**). To prepare a positively charged CPP-FP/Citrine complex, we increased the P/C ratio to 100, resulting in a complex with a hydrodynamic diameter of 880 nm and a zeta potential of 5.8 mV (**Fig. 2b**). Based on these results, different P/C ratios were used to optimize the size, PDI and surface charge for both the CPP-FP complexes.

**Figure 2.**
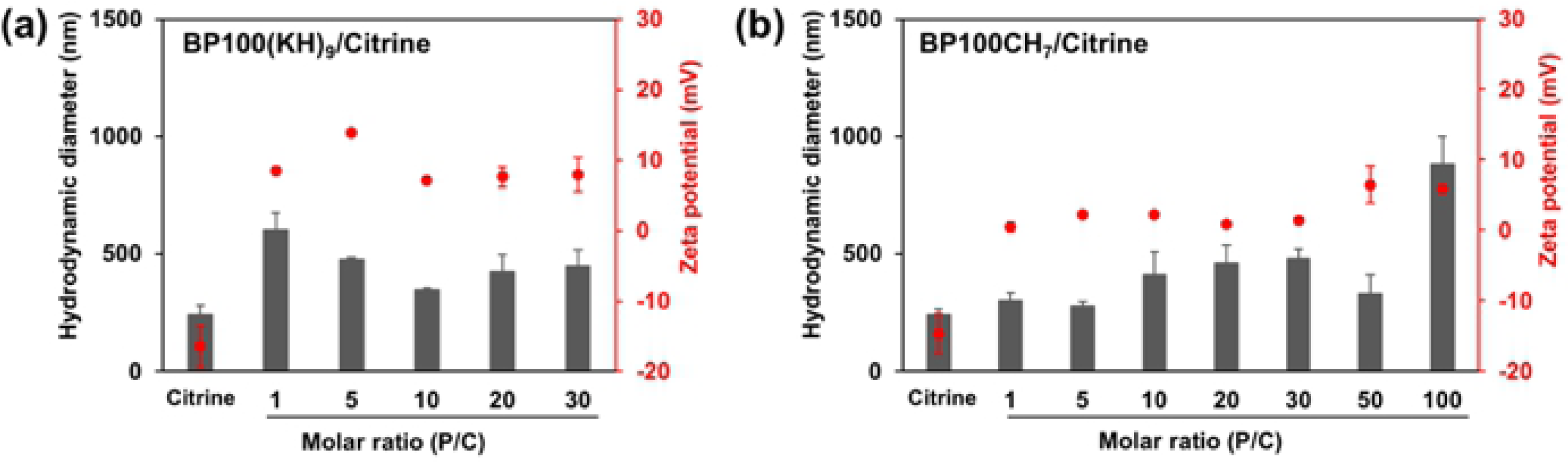
Size and zeta-potential values of CPP-FP/Citrine complexes prepared at different molar ratios. Data are presented as the means ± SD from triplicate tests.

### Regeneration of Rice Callus in Different Growth Stages

To assess the effects of different callus cell growth stages on CPP-FP/Citrine delivery efficiency, we used 5-day and 21-day rice callus cells as the recipient plant materials (**Fig. 3a- d**). To increase the surface area of the callus for efficient protein introduction with CPP-FPs, the callus was cut into pieces of various sizes. However, the damage caused by cutting callus can affect plant regeneration. We therefore tested the regeneration of the callus cells at different fragment sizes, from 0.5 to 6.0 mm in diameter. The smallest callus sizes shown in the red box could still generate plants (**Fig. 3e, f**); specifically, the 5-day callus could generate plants from callus fragments of 2.0 mm in size, while the 21-day callus showed excellent regeneration ability, even with callus fragments smaller than 0.5 mm. We thus prepared the 5-day callus with 2.0 mm diameter and the 21-day callus with 0.5 mm diameter for the rest of the experiments.

**Figure 3.**
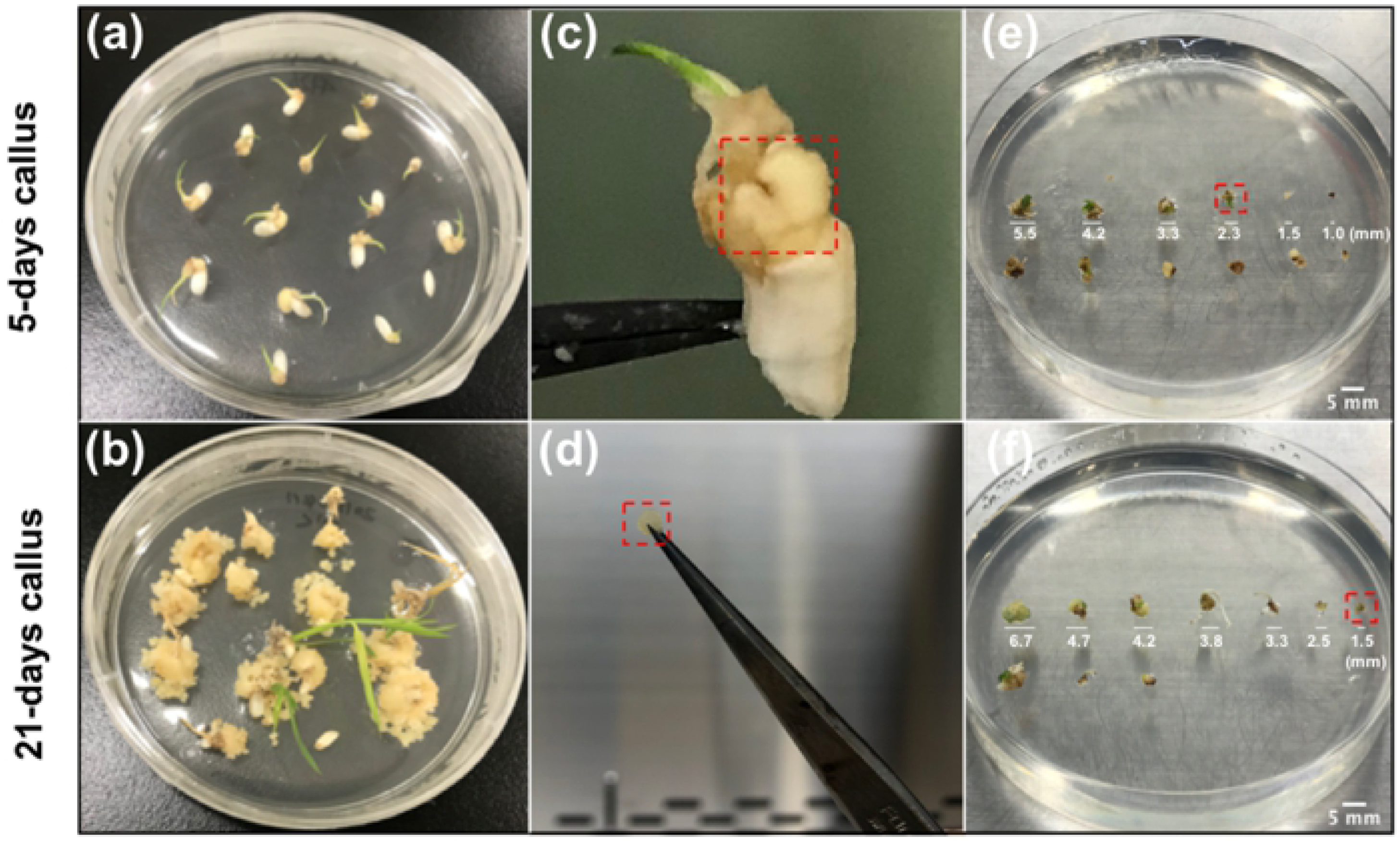
Regeneration test of the target cells for CPP-FP/Citrine complexes. Rice seeds were grown on N6D medium for 5 days or 21 days at 30°C with continuous light (a, b). Mature embryo-derived rice calli (c, d) were cut into small pieces of different sizes. The callus was then placed on regeneration medium and cultured under the same conditions for one week (e, f). The smallest calli used for plant generation are marked with red boxes.

### Selection of CPP-FP for Citrine Delivery into Rice Callus

To clarify the effect of CPP-FP on Citrine delivery efficiency into rice callus, two types of complex for each CPP-FP were investigated quantitatively. One had a smaller diameter with a low positive charge (P/C ratio of 1 for BP100(KH)_9_ and 5 for BP100CH_7_), and the other had a larger diameter with a high positive charge (P/C ratio of 10 for BP100(KH)_9_ and 100 for BP100CH_7_). Images were captured for observation of Citrine fluorescence signals every 24 hours by CLSM. Using the 5-day callus cells, the Citrine fluorescence inside the cells was first observed at 48 hours after infiltration with BP100(KH)_9_ at a P/C ratio of 10. The fluorescence signals were maintained up to 72 hours (**Fig. S1a**). Likewise, Citrine fluorescence inside the cells was observed using either the BP100(KH)_9_ complex at a P/C ratio of 1 or the BP100CH_7_ complex at a P/C ratio of 100 after 72 hours (**Fig. S1a**). Using 21-day rice callus, Citrine fluorescence was observed only for the BP100(KH)_9_ complex prepared at a P/C ratio of 10 after 48 hours (**Fig. S1b**). In addition, infiltration with Citrine showed fluorescence only in the intercellular spaces, which are the shared spaces between cells, at 72 hours (**Fig. S1a-b**). The negative control with an infiltration of Milli-Q water showed no fluorescence in any test. These results indicated that BP100(KH)_9_ and BP100CH_7_ are capable of delivering Citrine protein into rice callus.

### Protein Delivery Efficiency

To evaluate penetration efficiency at different recipient cell growth stages, we compared the Citrine fluorescence intensity per cell area in the 5-day and 21-day rice callus (**Fig. 4a-e**). The 5-day and 21-day calli were infiltrated with either the BP100(KH)_9_/Citrine complex at a P/C of 10 or the BP100CH_7_/Citrine complex at a P/C of 100. After 72 hours of incubation on N6D medium, the Citrine fluorescence intensity was quantified by CLSM for each cell. The average values and standard deviations from ten tests were calculated (**Fig. 4f**). The delivery efficiency of the BP100(KH)_9_/Citrine complex into the 5-day rice callus cell was approximately twice as high as that of the 21-day callus, and the difference was significant. Moreover, the BP100(KH)_9/_Citrine and BP100CH_7_/Citrine complexes showed similar fluorescence intensity values postinfiltration in 5-day callus cells, suggesting that, in comparison to the 21-day callus cells, the 5-day callus cells were better able to internalize the foreign cargo via CPP-mediated transmission. Furthermore, BP100(KH)_9_ and BP100CH_7_ were similar in delivery efficiency of Citrine protein into 5-day rice callus cells.

**Figure 4.**
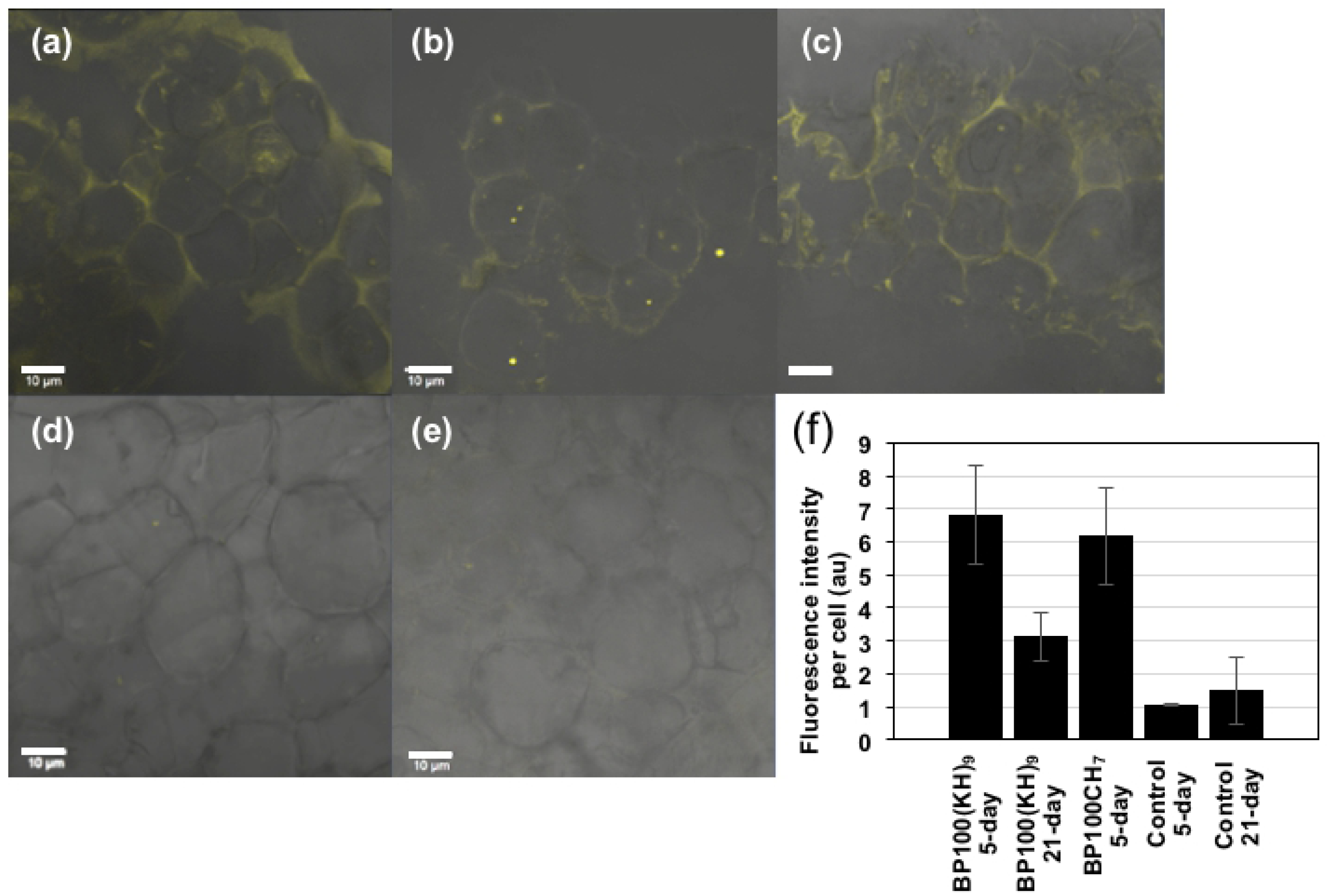
Quantification of Citrine fluorescence per cell. The single cell area was randomly selected, and their fluorescence intensity was calculated by CLSM. The results of Citrine delivery by BP100(KH)_9_ (a), BP100CH_7_ (c), and Milli-Q water (5) into the 5-day callus cells and of BP100(KH)_9_ (b) and Milli-Q water (e) into 21-day callus cells are shown. The SD indicates the standard deviations of ten analyses. The scale bar is 10 μm.

### Citrine Distribution in Callus Cells

To confirm the distribution of Citrine in callus cells, BP100(KH)_9_/Citrine complex at a P/C ratio of 10, BP100CH_7_/Citrine complex at a P/C ratio of 100, Citrine only, and Milli-Q water were infiltrated into 5-day rice callus cells. After 72 hours of incubation, the callus was stained with FM4-64 (**Fig. 5**) to mark the cell membrane. The merged CLSM images show overlapping fluorescence signals (orange) from Citrine and FM4-64 (**Fig. 5a-f**), suggesting that some Citrine proteins were localized on the cell membranes. Furthermore, the enlarged figures (**Fig. 5m, n**) showed yellow and orange spots inside the cells, indicating that Citrine had formed complexes with plasma membrane components. In addition, some red signals showed vesicles from the plasma membrane, whereas the Citrine-only and negative control (Milli-Q only) samples showed yellow signals in some cells, which might represent an endophyte, according to a previous report (**Fig. 5**, white arrows).

**Figure 5.**
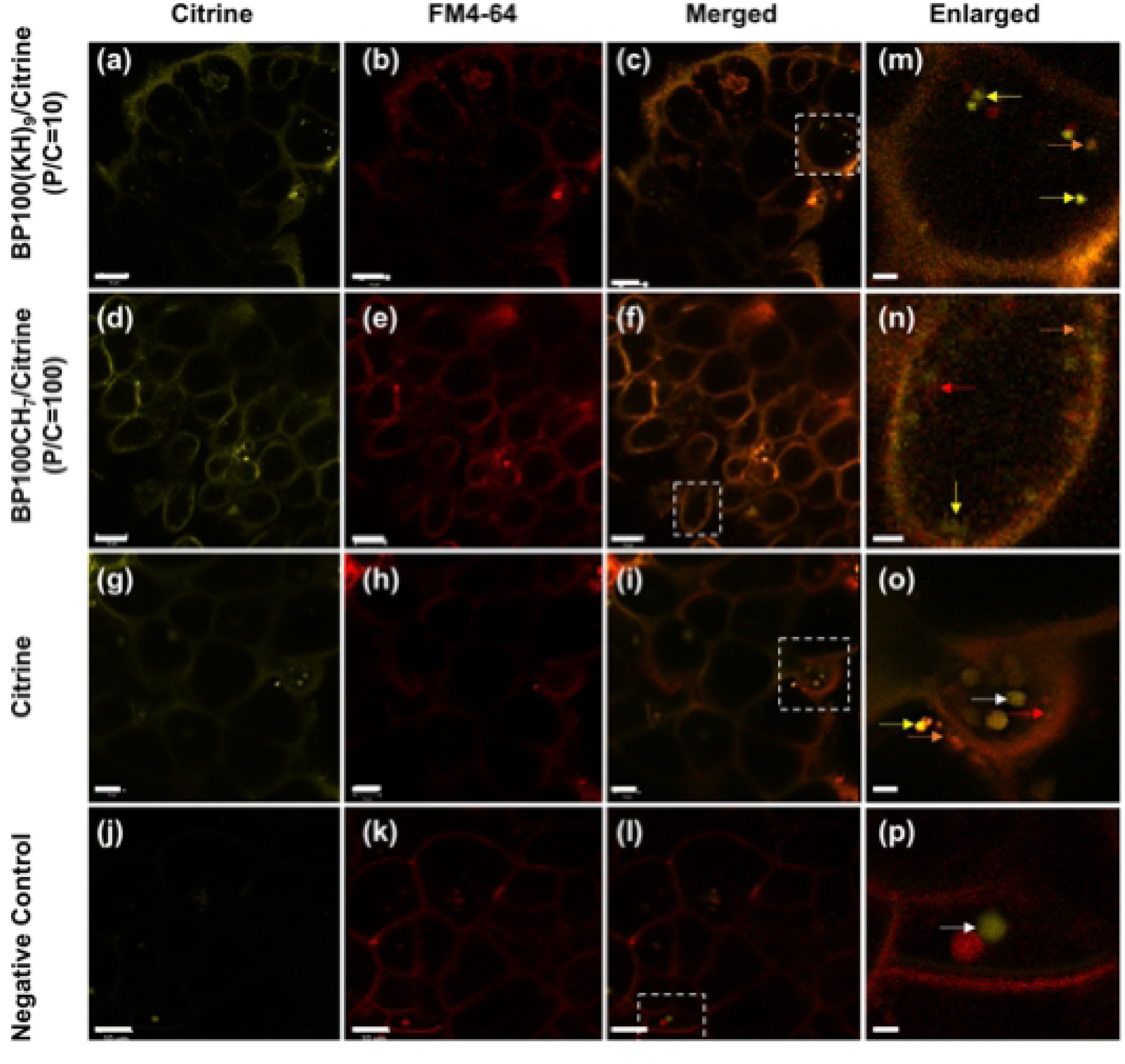
CLSM observation of intracellular distribution of BP100(KH)_9_/Citrine (g-i) and BP100CH_7_/Citrine (j-l) complexes by CLSM. The red color comes from FM4-64 dye, which was used for cell membrane staining. The yellow arrows indicate the Citrine position, the red arrows indicate the cell membrane stained with FM 4-64, and the orange arrows indicate a complex of Citrine and cell membrane fluorescence (m-p). The white arrows indicate symbiotic bacteria inside the cell (m-n). The negative control and Citrine represent samples infiltrated with Milli-Q water or Citrine only, respectively. The scale bar is 10 μm.

To further verify that the Citrine was delivered inside the cells, we generated 3D images from those CLSM images (**Fig. 6**). The results of BP100(KH)_9/_Citrine and BP100CH_7_/Citrine complexes showed that the yellow spots of Citrine signal occurred throughout the callus, and most of these spots were inside the cell membranes (**Fig. 6 a-b**, **Movie S1-2**). Consistent with **Fig. 5**, the red and orange spots were observed inside the cells, implying that Citrine was taken up with the plasma membrane into the cell via endocytosis. In contrast, the Citrine-only samples without CPP-FP showed a few yellow spots inside the cells (**Fig. 6c, Movie S3**), indicating that significantly less Citrine could be taken up without CPP-FP. To perform a confocal optical section analysis (**Fig. S2**), we randomly selected one yellow spot located in the *xy* plane inside the cells treated with the BP100(KH)_9_/Citrine or BP100CH_7_/Citrine complexes. The selected spots are displayed in the *xz* and *yz* planes. These results indicate that BP100(KH)_9_ and BP100CH_7_ can deliver Citrine proteins through the cell membrane into the cytoplasm.

**Figure 6.**
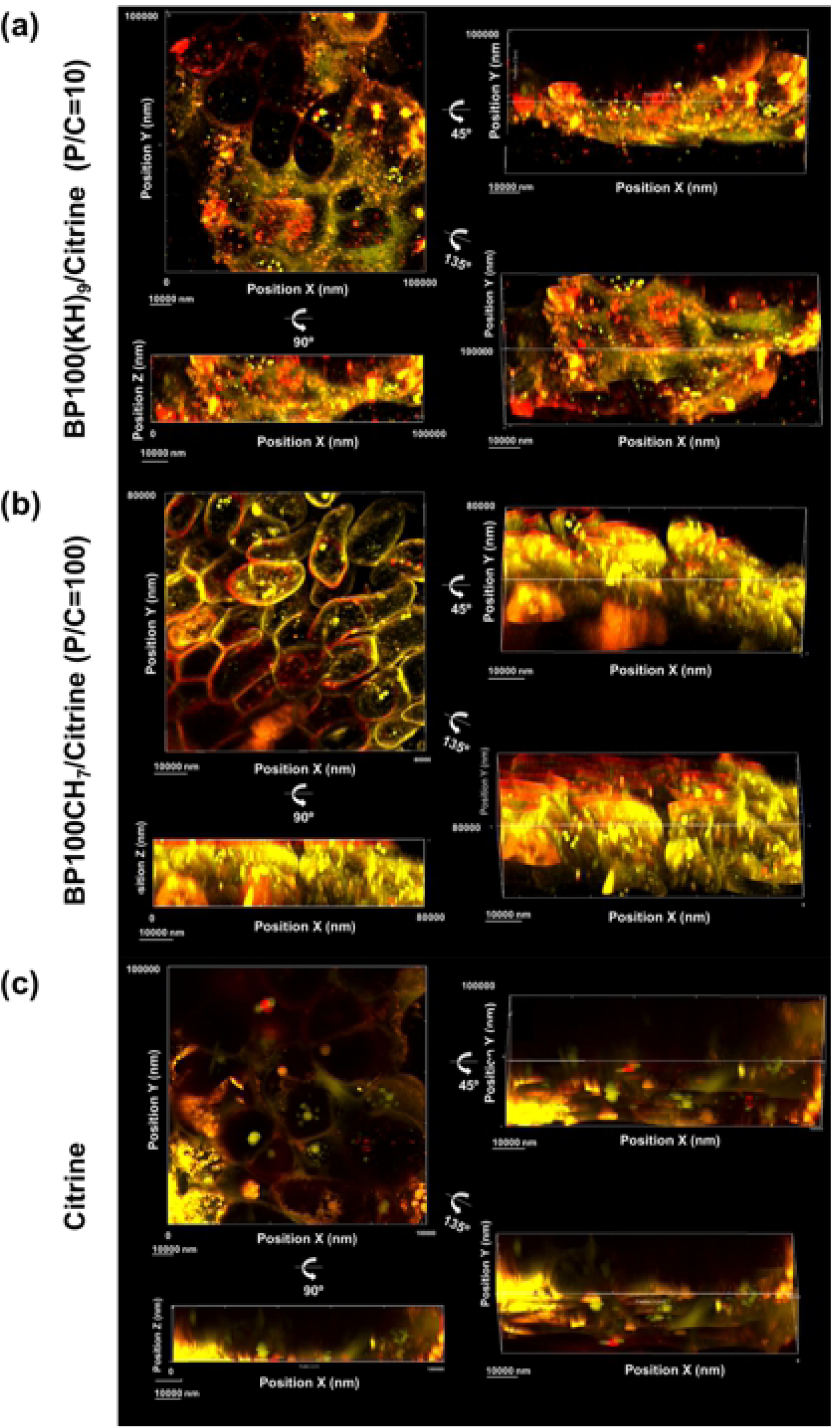
CLSM observation of intracellular Citrine delivery using BP100(KH)_9_, BP100CH_7_ and Citrine only. The yellow spots represent Citrine positions. The red spots represent the cell membrane stained by FM4-64. The orange spots represent the combination of peptide/Citrine and cell membrane fluorescence. The 3D structures were analyzed by Imaris software, and the pictures used are all from the CLSM Z-stack results.

### Quantification of Citrine Delivery by CPP-FP

We next quantified the Citrine extracted from the cell lysates by Western blot. The samples were prepared by infiltration of the BP100(KH)_9_/Citrine complex, the BP100CH_7_/Citrine complex and Citrine into 5-day callus cells. Samples with infiltration of Milli-Q water and Citrine only without vacuum and pressure treatment were also analyzed as controls (**Fig. 7a**). The results showed that, in addition to the BP100(KH)_9_/Citrine and BP100CH_7_/Citrine complexes, the infiltrated Citrine exhibited a Citrine protein band on the Western blot, even though both Milli-Q water and Citrine without vacuum and pressure treatment showed no Citrine. We quantified the band area intensities by comparison with the positive control (**Fig. 7b**). The amount of Citrine delivered without CPP-FP was approximately 0.024 ng/mg callus, whereas those delivered by BP100(KH)_9_ and BP100CH_7_ were approximately 0.042 and 0.043 ng/mg callus, respectively. Although the infiltrated Citrine exhibited a Citrine protein band on the Western blot, the infiltrated Citrine proteins were not taken up by the cells but were trapped in the intercellular spaces, according to the CLSM images (**Fig. 5 and Fig. 6**). Thus, the difference from the Citrine delivered without CPP-FP was considered to be the amount of Citrine introduced into the callus cells; namely, BP100(KH)_9_ and BP100CH_7_ delivered approximately 0.018 and 0.019 ng of Citrine per mg of rice callus.

**Figure 7.**
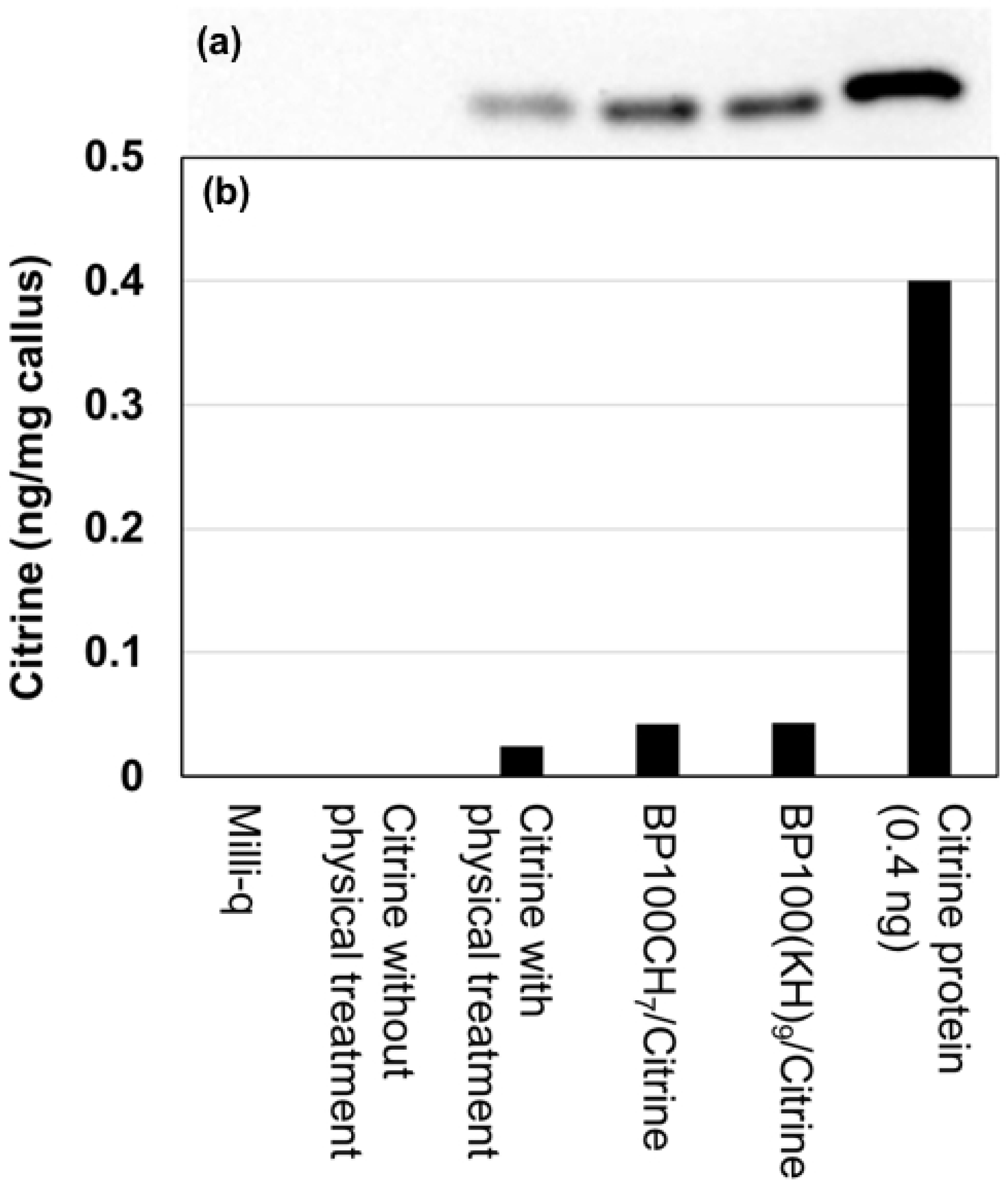
Western blot analysis of Citrine extracted from 5-day rice callus at 72 hours postinfiltration. (a) Western blot bands. (b) Quantitative analysis of those bands by ImageJ.

## Discussion

BP100(KH)_9_ is a cationic peptide synthesized by conjugating a cationic peptide sequence to a CPP, BP100 [20]. To form the ionic complex with the negatively charged protein, the CPP was fused to the cationic peptide composed of lysine and histidine, resulting in a fusion peptide, BP100(KH)_9_. This combination has been proven for high penetration efficiency of protein, DNA and RNA into plant cells [21-23]. BP100CH_7_ is another fusion peptide containing a cyclic pDNA-binding domain and was designed as a DNA carrier [16]. In this study, we used BP100CH_7_ for protein delivery; however, its protein binding mechanism has not yet been clarified. BP100CH_7_ may interact with Citrine protein via ionic interactions based on its zeta-potential measurements (**Fig. 2b**).

The cell membrane is a structural barrier that protects living cells from the surrounding environment. In general, electrostatic interaction with negatively charged phospholipids is considered an essential factor for CPP-mediated cargo delivery [10]. Hence, both final sizes and surface charges are thought to affect the penetration behavior of the CPP-FP/Citrine complex. In our results, the BP100(KH)_9_/Citrine complex at a P/C ratio of 10 and the BP100CH_7_/Citrine complex at a P/C ratio of 100 exhibited more rapid delivery than those with lower P/C ratios. However, both of these complexes showed relatively larger hydrodynamic diameters and highly positive surface charges, namely, 456 nm and 7.2 mV for BP100(KH)_9_/Citrine and 986 nm and 5.8 mV for BP100CH_7_/Citrine, than the other complexes tested, suggesting that the surface charges of the complexes were more important to the delivery efficiency than were the complex sizes (**Fig. 2, Table S1**).

Rice callus, which is induced from rice scutellum and cultured for 21 days, is widely used as the recipient in rice genetic modification by Agrobacterium-mediated transformation and is also used for CPP molecular delivery due to its strong regeneration ability [2,19]. However, transformation usually requires a 3-month or even longer experimental period to develop a transgenic plant. Notably, 5-day rice callus has been reported to show similar transformation efficiency to 21-day callus [5]. In the current study, we compared 5-day and 21-day rice calli as recipient cells in terms of protein delivery efficiency. In comparison to the 21-day callus, the 5-day rice callus exhibited a distinct advantage for CPP-mediated protein delivery due to its higher efficiency (**Fig. 4**). This effect could be because rice calli at different growth stages contain different lipids and metabolites that affect the cellular uptake [14].

The subcellular fluorescence signals from Citrine and FM4-64 demonstrated that both BP100(KH)_9_/Citrine and BP100CH_7_/Citrine could pass through the rice cell wall (**Fig. 5a-f**). This result agrees with the result shown in **Fig. 6a-b**. Compared to the CPP-FP/Citrine complexes, Citrine without CPP-FP showed few fluorescence signals inside the cells (**Fig. 5g-i, Fig. 6c**), and these fluorescence signals did not come from the cytoplasmic region but rather the intercellular spaces (**Fig. 5o, Fig. 6**). The Western blot results revealed that Citrine without CPP-FP can be introduced into rice callus by vacuum and pressure treatment (**Fig. 7c**). Considering the Western blot results and CLSM image analyses, we conclude that CPP-FP is necessary for the internalization of Citrine protein in rice callus cells via the plasma membrane.

In conclusion, this study is the first time that clear evidence has been provided of the utility of BP100(KH)_9_ and BP100CH_7_ for protein delivery into rice callus cells. Moreover, our results are the first to demonstrate that 5-day rice callus cells are suitable recipients for CPP-mediated protein delivery. The CPP-mediated Citrine delivery process is proposed to occur through the following steps. First, CPP-FP-Citrine complexes pass through the cell wall into the intercellular space of rice callus by a combination physical treatment of vacuum followed by pressure, and these complexes are then transferred into cells by the interaction between CPP and the lipid bilayer. Thus, the fluorescence signals of Citrine were detected both on the cell membrane and inside the cells. By contrast, the Citrine without CPP-FP was delivered in the same way; however, the Citrine accumulated in the intracellular space and could not be internalized without CPP. This protein delivery system illustrates the possibility of DNA-free genetic modifications in higher plant cells. In our ongoing work, we expect to use this protein delivery system for targeted epigenetic genome editing via Cas9/gRNA delivery to explore novel trait improvement in rice. This method could be widely applicable for producing genome-edited crop plants and has the potential to be commercialized.

## Acknowledgments

The present study was financially supported by BASF SE and grants from the Japan Science and Technology Agency Exploratory Research for Advanced Technology (JST-ERATO; Grant No. JPMJER1602).

## Supporting Information

**Figure S1. Time-course analyses of Citrine delivery by BP100(KH)**_**9**_ **and BP100CH**_**7**_ **at different P/C molar ratios.** Five-day (a) and 21-day rice calli (b) were used as recipient cells. The negative control represents samples treated with Milli-Q water only. Scale bar: 10 μm.

**Figure S2. Confocal optical section analysis (xy, xz, and yz planes) of Citrine delivery by BP100CH**_**7**_**, BP100(KH)**_**9**_ **and Citrine without additional peptides.** The green, red and dark blue are the contour lines for the X, Y, and Z directions. The arrows indicate the yellow spots representing the Citrine positions in different directions.

**Table S1. Characterization of BP100(KH)**_**9**_**/Citrine and BP100CH**_**7**_**/Citrine complexes prepared at various P/C ratios.**

